# Rational design of tertiary coordination sphere of a heme-based sensor for two-orders enhanced oxygen affinity

**DOI:** 10.1101/2025.05.27.656425

**Authors:** Anoop Rama Damodaran, Eaindra Yee, Rahul L. Khade, Murphi Williams, Elizabeth A. Apiche, R. Hunter Wilson, Edward Hoey, Grant Larson, Ke Shi, Hideki Aihara, Yong Zhang, Ambika Bhagi-Damodaran

## Abstract

Biological O_2_ sensing is crucial for diverse physiological functions across all forms of life. Heme-containing proteins achieve this by binding O_2_ to their iron center and have been found to display O_2_ affinities spanning several orders of magnitude. Despite decades of investigation into the structure and function of heme-based O_2_ sensors, the molecular mechanisms that enable the tuning of O_2_ affinity to match specific physiological roles remain unclear. Here, we utilize the O_2_ sensing mycobacterial DosS protein as a model system to explore the role of heme iron’s tertiary coordination sphere in controlling its O_2_ affinity. By rationally and systematically modifying the tertiary coordination sphere to promote the formation of a Trp-Tyr-Asn H-bond triad within the heme’s distal pocket, we have enhanced the O_2_ affinity of WT DosS by over 150-fold. The rationally designed DosS exhibited a *K*_d_ value of 3 ± 1 nM, compared to 460 ± 80 nM for WT DosS. Employing a combination of structural, biochemical, spectroscopic, and computational studies, our analysis of WT and designed DosS variants highlights how the interplay between distal H-bond networks and heme-pocket electrostatics drives large differences in their O_2_ sensing capabilities. Ultimately, our work shows how metalloenzymes can dramatically alter their sensitivity to diatomic signaling molecules by tuning the tertiary coordination sphere, broadly impacting how we understand related biological sensing and signaling.

## Introduction

O_2_ is a vital metabolite and cells possess intricate molecular pathways for precisely sensing O_2_ concentrations.^1^ Cellular O_2_ levels are primarily detected by iron-containing metalloproteins that employ heme, non-heme, or iron-sulfur cofactors.^2^ Among these, heme-based O_2_ sensors are the most extensively studied and are found to exhibit O_2_ affinity values spanning six orders of magnitude.^3^ In heme-based O_2_ sensors, O_2_ binds to heme iron in an end-on fashion with a partial negative charge on its atoms which can be stabilized by H-bonds from residues within the secondary coordination sphere (SCS).^4^ For instance, tyrosine with its significant H-bond donating capability (pKa = 10) is a frequently utilized residue in the SCS of heme-based bacterial O_2_ sensors (**Fig. 1a, Table S1**). The absence of such an H-bond in the O_2_-sensing enzyme FixL results in its significantly weakened O_2_ affinity of 130 μM.^5^ In the globin-coupled heme-based O_2_ sensor YddV featuring a tyrosine residue (Y43) in the SCS, its Y43A and Y43L mutants display dramatically low O_2_ affinities to the point that their Fe^II^-O_2_ complexes have not been spectrally detected.^6^ On the other hand, the introduction of two secondary coordination H-bonding residues (I145Y and I149N) into sGC confers upon this otherwise O_2_-inert enzyme the ability to bind and interact with O_2_.^7^ In all, these studies highlight the crucial role of SCS residues in determining the O_2_ sensing capability of heme-based sensors.

**Fig. 1:**
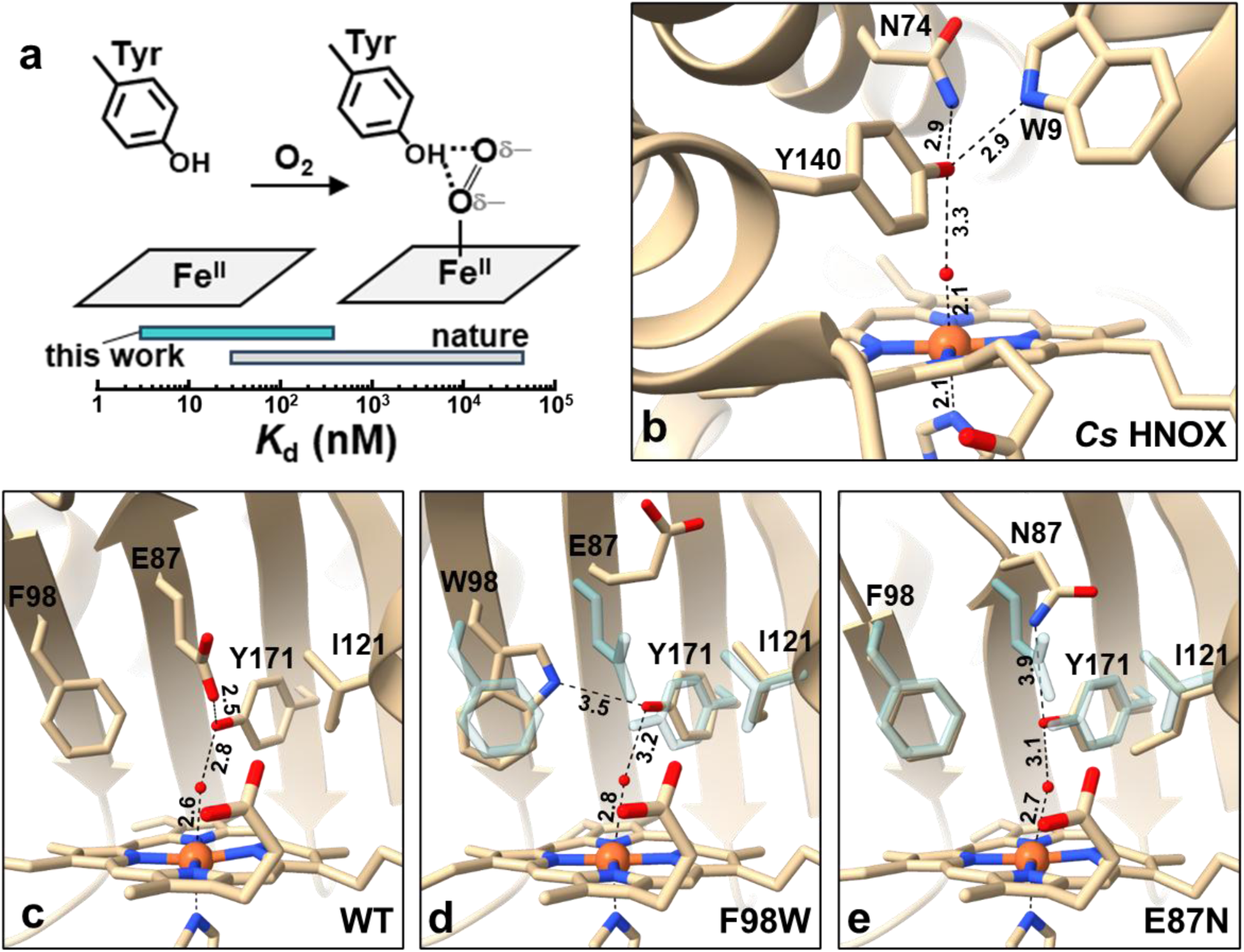
a) Schematic shows O_2_ binding to ferrous heme. The partial negative charge on iron-bound O_2_ is stabilized via H-bonding to SCS tyrosine residue. The graph shows the range of Kd value spanned by heme-based O_2_ sensors that feature tyrosine in their SCS (in gray). The cyan bar represents the Kd value spanned by DosS sensors designed in this work that feature tyrosine in their SCS. b) Crystal structure of ferric form of Cs H-NOX that zooms into its distal heme pocket. H-bonding distances are shown via dashed lines. c) Crystal structure of ferric form of WT, d) F98W, and e) E87N GAF-A DosS that zooms into their distal heme pocket. H-bonding distances are shown via dashed lines. Blue structure in d-e show corresponding SCS/TCS residues in WT GAF-A DosS overlaid.

While the importance of H-bonding SCS residues in modulating O_2_ binding in heme-based sensors is well recognized, the molecular mechanisms that drive large variations in O_2_ affinities among heme-based sensors sharing the same residue in their SCS remain unclear. For instance, heme proteins with tyrosine in their SCS exhibit *K*_d_ values that span three orders of magnitude (**Table S1**). These include the mycobacterial heme-based sensor DosS,^8,9^ featuring Y171 in its SCS and an O_2_ *K*_d_ of ∼460 nM^10^, while H-NOX from *C. subterraneus* with Y140 in its SCS^11–13^ exhibits O_2_ affinities that an order-of-magnitude stronger. Notably, the Y140 SCS residue of *Cs* H-NOX is held in position by an H-bond triad with H-bond donating tertiary coordination residues, aspargine (N74) and tryptophan (W9) (**Fig. 1b**). Mutation of W9 to phenylalanine weakens its O_2_ affinity by ∼3.4-fold^13^, suggesting that residues beyond the SCS could play a role in the stabilization of the Fe^II^-O_2_ species. In contrast to *Cs* H-NOX, the SCS Y171 of DosS is held in position by an H-bond withdrawing tertiary coordination glutamate (E87) residue (**Fig. 1c**). Differences in these tertiary coordination sphere (TCS) residues and the nature of the H-bonding networks they form with SCS residues could provide additional mechanisms for tuning the O_2_ affinity of heme-based O_2_ sensors.

In this work, we focus on engineering the TCS of *M. tuberculosis* DosS, an extensively characterized heme-based O_2_ sensor, to control and enhance its O_2_ binding affinity. WT DosS features an H-bond-accepting E87 residue in its TCS capable of forming an H-bond with the Y171 phenol sidechain. We introduced systematic mutations into the TCS to promote the formation of an H-bond triad akin to that in *Cs* H-NOX. By mutating E87 to asparagine to generate the E87N DosS mutant and subsequently incorporating F98W and I121L mutations to generate the NWL DosS mutant, we have systematically strengthened O_2_ binding affinity from 460 ± 80 nM for WT DosS to 43 ± 13 nM for E87N and ultimately to 3 ± 1 nM for NWL DosS – an enhancement of over two orders of magnitude. Employing a combination of structural, biochemical, spectroscopic, and computational studies, our analysis of WT and designed DosS variants highlights how the interplay between distal H-bond networks and heme-pocket electrostatics drives dramatic differences in their O_2_ sensing capabilities. At the same time, we demonstrate the potential of computation-guided protein design for engineering the TCS of metalloprotein sensors to achieve desired O_2_ sensitivity. Overall, our studies provide insights into longstanding questions regarding the interplay between SCS and TCS residues of metalloproteins in determining their sensitivity to diatomic signaling molecules.

## Results and Discussion

We begin by examining the crystal structure of *Cs* H-NOX which exhibits the tightest O_2_ affinity among heme-based O_2_ sensors that feature a tyrosine at its SCS (**Table S1**). Focusing on the distal heme pocket, we observe that *Cs* H-NOX positions its Y140 residue in close proximity to H-bond donating W9 and N74 residues to form a distinctive H-bond triad motif (**Fig. 1b**). To understand the role of such extended H-bond networks in controlling the O_2_ affinity of heme-based sensors, we decided to incorporate elements of this triad into DosS. We note that the heme-containing GAF-A domain of DosS is well-suited for structural investigations via X-ray crystallography^14,15^ and provides a viable platform to explore the structural implications of these designed mutations. Furthermore, its distinct three-tiered α/β fold (different from *Cs* H-NOX’s predominantly helical fold) allows us to probe the impact of distal H-bond networks on O_2_ affinity in a topologically distinct environment.

We expressed and purified WT GAF-A DosS along with single mutant variants F98W and E87N for structural investigations, wherein elements of the H-bond triad are individually incorporated in the TCS of heme iron (**Fig. S1-S2**). The ferric form of the purified proteins were characterized with UV-Vis spectroscopy and they all exhibit a sharp Soret peak at 406 nm and a broad visible transition in the 500-650 nm region indicative^9,16^ of a well-folded heme containing protein (**Fig. S3**). We have successfully crystallized WT, F98W and E87N GAF-A DosS in their ferric form (see method for details on protein crystallization). Synchrotron X-ray diffraction studies of these crystals reveal that all GAF-A DosS variants crystallize as dimers in the asymmetric unit of crystals with orthorhombic space group (**Table S2**). Zooming into the distal heme pocket of WT GAF-A DosS (**Fig. 1c**), we see that the ferric heme is coordinated to an aqua ligand which is within H-bonding distance (2.8 Å) of Y171’s hydroxyl group. Focusing on the TCS, we note that the H-bond withdrawing carboxylate side chain of E89 is also within H-bonding distance (2.5 Å) from Y171’s hydroxyl group. Next, we examine single mutants, F98W and E87N (**Figs. 1d-1e**), both of which display excellent overall structural conservation and overlay with WT (RMSD of 0.68 Å for F98W and 0.56 Å for E87N). They show subtle changes in the positioning and sidechain orientations of crucial SCS and TCS residues distal to heme as compared to WT (selected WT residues overlaid in transparent blue, **Figs. 1d-1e**). For the F98W mutant, the phenyl ring of W98’s indole sidechain occupies a position analogous to F98 in WT. However, the added steric bulk from W98’s pyrrole ring causes E87’s carboxylate side chain to flip upward and reorient away from the heme center. Furthermore, W98’s indole N1 is positioned within H-bond distance (3.5 Å) from Y171’s hydroxyl group. Here, Y171’s hydroxyl group is located ∼1 Å away from its WT location to facilitate this H-bond with W98. We specifically pursued the E87N mutant (over isosteric glutamine) to both mimic the Asn-Tyr H-bond found in *Cs* H-NOX and to mitigate potential steric clashes when combined with the F98W mutation. For the E87N mutant, the amide sidechain of N87 is oriented with its amine group facing Y171’s hydroxyl and forms a weak H-bond (3.9 Å) (**Fig. 1e**). Combining this E87N mutation with F98W which repositions Y171 farther from the iron center, could create conditions conducive to forming an H-bond triad. Overall, our crystallographic studies of DosS’s GAF-A domain suggest that both F98W and E87N are capable of H-bonding to Y171, and in combination could potentially form distal H-bond networks analogous to *Cs* H-NOX.

In order to mitigate challenges related to the steric bulk of W98, an additional I121L mutation was included in the variant that combines E87N and F98W to yield the NWL DosS variant. Despite extended efforts, crystallization of NWL GAF-A DosS proved unsuccessful. Consequently, we employed molecular dynamics^17,18^ (MD) simulations of the O_2_-bound form of NWL GAF-A DosS to investigate potential distal H-bond networks. These studies reveal that NWL DosS is able to sample configurations with extensive H-bonding between O_2_, SCS and designed TCS residues (**Fig. 2a**). To experimentally explore the role of distal H-bond networks, we selected WT, E87N, and NWL DosS as a useful model system. This set of proteins systematically varies the H-bond environment influencing the heme-bound O_2_ and the SCS Y171 residue. More specifically, WT DosS features an H-bond withdrawing E87 residue in the TCS, E87N introduces a potential H-bond donor from N87’s amine group to Y171, while NWL DosS uniquely features two H-bond donors (W98 and N87) interacting with Y171. We expressed and purified full-length versions of DosS mutants (**Fig. S4-S5**) due to their enhanced robustness against autooxidation compared to isolated GAF-A domains. We note that *Cs* H-NOX is known to dissociate its proximal heme-bound histidine upon NO binding.^19^ This effect is attributable to its distal H-bond triad stabilizing the Fe-NO bond,^20^ which in turn weakens the Fe-His bond via trans effect. To investigate if similar effects occur in DosS variants, we incubated their ferrous forms with >10 equivalents of NO in an anaerobic setting. Both WT and E87N DosS formed a stable NO-bound species with a Soret maximum at 419 nm, typical of a 6-coordinate (*6c*) NO/His-bound heme complex (**Fig. 2b, S6**). In contrast, NWL DosS yielded a mixture of *6c* and *5c* Fe-NO species, indicated by Soret bands at 419 nm and 399 nm,^20^ respectively (**Fig. 2c**). The observed proximal histidine dissociation in NWL DosS resembles the behavior of *Cs* H-NOX with NO and strongly suggests the presence of a distal H-bond network in NWL FL DosS comparable to that found in *Cs* H-NOX.

**Fig. 2:**
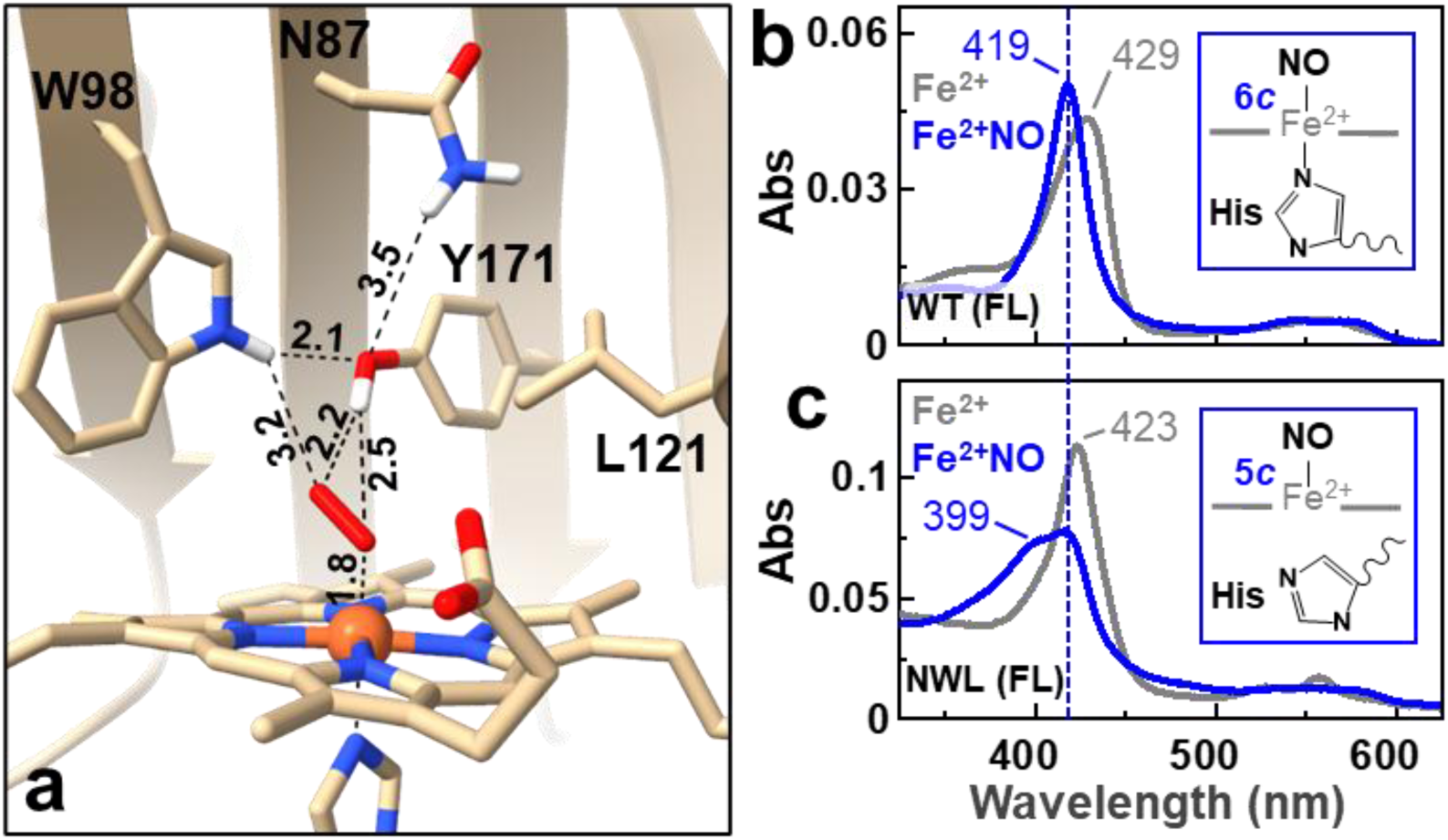
a) A distal heme pocket zoomed-in snapshot from MD simulations of O_2_-bound NWL GAF-A DosS. H-bonding distances are shown via dashed lines. UV-Vis spectroscopic studies showing ferrous forms of b) WT and c) NWL DosS (in gray) binding to NO (in blue). Inset in b) shows the structure of a 6c NO/His bound ferrous heme and c) shows the structure of a 5c NO bound ferrous heme with proximal histidine dissociated from the heme iron.

Next, to assess the impact of designed TCS mutations on O_2_ binding, we performed equilibrium binding affinity measurements by titrating O_2_ to the reduced ferrous form of DosS variants. We used UV-Vis spectrometry to determine the O_2_-bound fraction via Soret band shifts (**Fig. 3a-b** and **S7-8**), while simultaneously employing an O_2_ optode to measure the free dissolved O_2_ concentration at each titration step.^21^ A Hill’s fit with n=1 for the O_2_-bound fraction versus the corresponding free dissolved O_2_ yielded the *K*_d_ value. We measured a *K*_d_ of 43 ± 13 nM (purple curve, **Fig. 3c**) for O_2_ binding to E87N DosS, which is over 10-fold tighter than WT DosS (*K*_d_ = 460 ± 80 nM^10^ from dark blue curve, **Fig. 3c**). NWL DosS exhibited a further 14-fold enhancement in O_2_ binding affinity (*K*_d_ = 3 ± 1 nM from maroon curve, **Fig. 3c**) over E87N. The O_2_ affinity of our designed NWL DosS mutant is an order-of-magnitude stronger than any previously reported WT heme-based O_2_ sensor featuring tyrosine in its SCS (**Fig. 1a**). Collectively, these measurements underscore the critical role of TCS residues in determining the O_2_ sensitivity of heme-based sensors.

**Fig. 3:**
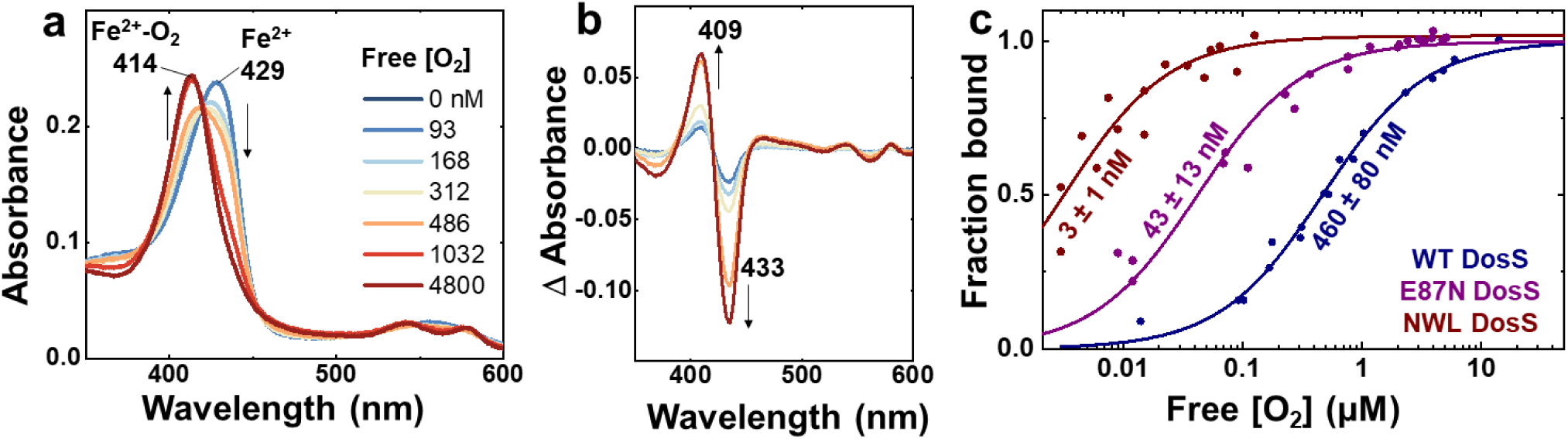
a) UV–Vis spectral changes in WT DosS upon binding O_2_ at various free O_2_ concentrations measured using the optode. b) Difference spectra showing spectral changes when WT DosS binds O_2_. c) O_2_ affinity plots for WT (dark blue), E87N (purple), and NWL (maroon) DosS proteins (n = 3).

To understand the molecular basis of the 150-fold enhanced O_2_ affinity of NWL DosS over WT DosS, we employed computational studies including density functional theory^16^ (DFT) and MD. DFT calculations were performed on heme-O_2_ adducts of WT and NWL DosS along with their primary, secondary and tertiary coordination residues. We find that incorporating the H-bond triad in NWL DosS dramatically alters heme pocket electrostatics which has significant implications for O_2_ binding. The negatively charged E87 residue in WT DosS creates an overall negative electrostatic potential near the heme pocket (**Fig. 4a**). Since O_2_ binds to heme with a partial negative charge on its atoms,^22^ this negative potential hinders O_2_ binding in WT DosS. In contrast, the mutation of E87 to a neutral aspargine in NWL DosS results in a more neutral heme pocket (**Fig. 4b**) that favors O_2_ binding. While we initially anticipated H-bond donation from N87 to Y171 to be a major contributing factor, MD distance distributions reveal that these interactions are sampled for less than 10% of the simulation time and correspond to weak H-bonds (**Fig. 4c**). In combination with DFT studies, these observations suggest that the contribution of E87N mutation is mostly electrostatic in nature. Focusing on the F98W mutation in NWL DosS, MD distance distributions (**Fig. 4d**) reveal strong H-bond donation from W98’s indole N1 to Y171, which in turn strengthens Y171’s H-bonding to the heme-bound O_2_. Moreover, F98W also samples direct H-bonding distance with heme-bound O_2_ for 30% of the simulation (**Fig. 4e**), suggesting it can stabilize O_2_ through direct H-bond donation. Ultimately, these studies highlight a complex interplay of pocket electrostatics and H-bond donation from SCS and TCS residues that can be used to tune the sensitivity of heme-based O_2_ sensors.

**Fig. 4:**
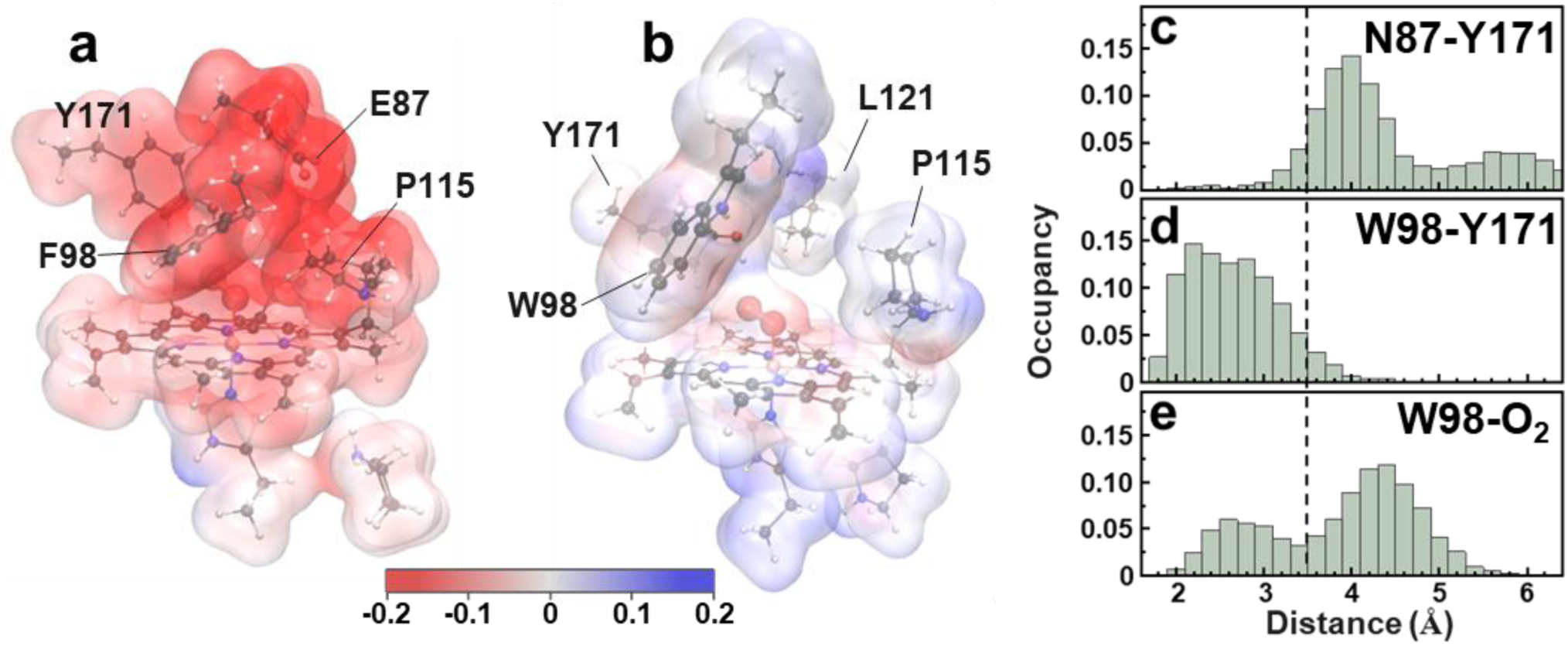
a) Electrostatic potential maps of a) WT and b) NWL DosS obtained using their DFT simulated heme bound to O_2_ structures. Histogram depicting distances sampled between c) N87’s amine hydrogens and Y171’s hydroxyl oxygen, d) W98’s N1 hydrogen and Y171’s hydroxyl oxygen and e) W98’s N1 hydrogen and distal oxygen atom of heme-bound O_2_ in MD simulated structure of O_2_-bound NWL DosS. Dashed line in c-e represents cut-off of H-bonding distances at 3.5 angstrom.

## Conclusion

Heme-based sensors are vital in the redox sensing and signaling pathways of numerous microorganisms and exhibit a remarkable million-fold dynamic range across biological systems. These sensors precisely modulate their O_2_ affinity by adapting their protein architecture. Heretofore, the wide variety of H-bond donating secondary SCS residues (e.g., tyrosine, histidine, arginine, threonine) in these sensors were thought to be the primary determinants of their O_2_ affinity. Here, we demonstrate how TCS and SCS cooperatively determine the O_2_ affinity of heme-based sensors. Through rational design of the TCS in DosS and the incorporation of an extensive H-bond network within its distal heme pocket, we enhanced its O_2_ affinity from 460 ± 80 nM for WT to 3 ± 1 nM for the designed NWL DosS variant. Notably, NWL DosS exhibits an order-of-magnitude stronger O_2_ affinity than *Cs* H-NOX, despite the latter also possessing a strong H-bond network in its distal pocket. This could be due to the presence of a distorted heme in *Cs* H-NOX^11^, which hinders d*π*-p*π* back-bonding to heme-bound O_2_, which in turn weakens its Fe-O bond. We believe this work serves as a foundational step for researchers to consider TCS interactions as a key factor in controlling substrate binding and molecular recognition. Beyond O_2_ sensing, heme-based sensors are integral to recognizing physiological levels of other biologically relevant redox-active stimuli such as NO, CO, and H_2_S. From a redox sensing and signaling perspective, this study provides clues to how primary, secondary, and tertiary coordination sphere residues are strategically designed in proteins to achieve desired sensitivity to diverse redox stimuli. Developing this understanding is crucial for elucidating the molecular basis of signal transduction in biology. Furthermore, from an enzyme design perspective, this work underscores the immense potential of rational and computational protein design strategies in precisely controlling the dynamic range of a wide array of metalloprotein-based sensors.

## Acknowledgement

ARD, EY, MW, EAA, EH, GL, and ABD acknowledge the support of NIH NIGMS grant #R35GM138277 for funding this work. KS and HA were supported by R35GM118047. RHW was supported by NIH Chemical Biology Training Grant #T32GM132029. RK and YZ were supported by NIH grant R15GM085774. The X-ray diffraction data were collected at the Northeastern Collaborative Access Team (NECAT), SSRL 12-2 and NSLSII 17-ID-1 beamlines. NECAT is funded by the NIH (P30 GM124165). The Pilatus 6M detector on 24-ID-C beamline is funded by a NIH-ORIP HEI grant (S10 RR029205). Use of the Stanford Synchrotron Radiation Light source, SLAC National Accelerator Laboratory, is supported by the U.S. Department of Energy, Office of Science, Office of Basic Energy Sciences under Contract No. DE-AC02-76SF00515. The SSRL Structural Molecular Biology Program is supported by the DOE Office of Biological and Environmental Research, and by the NIH/NIGMS (P30GM133894). The Center for Bio-Molecular Structure (CBMS) is primarily supported by the NIH-NIGMS through a Center Core P30 Grant (P30GM133893), and by the DOE Office of Biological and Environmental Research (KP1607011). NSLS-II is a U.S.DOE Office of Science User Facility operated under Contract No. DE-SC0012704. This publication resulted from the data collected using beamtime obtained through NECAT BAG proposal # 311950.

## Author Contributions

ARD and ABD designed and supervised this study. ARD developed and performed all the biochemical and spectroscopic measurements. EY expressed and purified DosS proteins and conducted hemochromogen assays on them. EH purified and crystallized WT GAF-A DosS, MW purified and crystallized F98W GAF-A DosS, EAA purified and crystallized E87N GAF-A DosS. KS and HA conducted diffraction studies on DosS GAF-A crystals and solved the diffraction data. RHW performed MD simulations. GWL developed His-tag cleavage protocols. RK and YZ performed DFT calculations and analysis. ARD and ABD wrote the manuscript with contributions from all authors. All authors have given approval to the final version of the manuscript.

## Notes

### Competing Interest Statement

The authors have declared no competing interest.

### Summary of Updates

changes in main text and figures

