## Supplementary material for "Rational design of tertiary coordination sphere of a heme-based sensor for two-orders enhanced oxygen affinity": SI

**Materials and Methods:**

**Expression and purification of DosS GAF-A variants:**

A pet28a(+) plasmid harboring WT DosS GAF-A gene was custom synthesized and procured from Genscript. Site-directed mutagenesis was performed to obtain E87N and F98W GAF-A mutants. The plasmids were each co-transformed with GroES/EL plasmid into BL21-Gold (DE3) competent cells (Agilent) per commercial protocol. The cells were then plated onto an LB-agar plate supplemented with ampicillin (100 µg/mL) and chloramphenicol (37 µg/mL) and grown overnight at 37°C. A single colony was picked from the plate, inoculated into 2XYT medium supplemented with the same antibiotic concentrations, and grown overnight at 37°C with shaking (200 rpm). 10 ml of the overnight culture was used to inoculate 1 L 2XYT medium containing 100 µg/mL ampicillin and 37 µg/mL chloramphenicol. The cells were then grown to optical density, $OD_{600}$, of 0.6-0.8 before adding 30 mg/L hemin and being induced with 1 mM IPTG at 18°C for 24 hours. After overexpression, the cells were harvested by centrifugation at 8,000 rpm at 4°C, flash-frozen, and stored at -20°C until usage.

For purification, the harvested cell pellet was resuspended in lysis buffer (50 mM $NaH_2PO_4$, pH 7.5, 250 mM NaCl, 10% glycerol, 1% Triton X-100, 0.1 mg/mL DNase, 0.1 mg/mL RNase, 1 mg/mL lysozyme, 1 tablet/50 mL Pierce™ Protease Inhibitor), sonicated, centrifuged (20,000 rpm, 4°C), and filtered (using MCE Membrane Filter, 0.22 µm) before loaded onto 5-mL Histrap FF column on an AKTA Start protein purification system. After sample application, the unbound proteins were washed out with 10 CV of IMAC buffer A (50 mM $NaH_2PO_4$, pH 7.5, 500 mM NaCl, 20 mM imidazole, 10% glycerol). GAF-A proteins were eluted using a two-step gradient method: 10% IMAC buffer B (50 mM $NaH_2PO_4$, pH 7.5, 500 mM NaCl, 400 mM imidazole, 10% glycerol) for 7 CV, then 100% buffer B for 5 CV. Both the wash out unbound and the elution steps were performed at the flow rate of 2 mL/min. The protein fractions from the last elution step were pooled, buffer-exchanged into storage buffer (50 mM tris-HCl, pH 7.6, 100 mM NaCl, 5% glycerol), and incubated with TEV protease (1:20 w/w TEV:protein) at 4°C overnight to cleave off the His-affinity tags from GAF-A.

The His-tag removed GAF-A proteins were again run on the 5-mL Histrap FF column on AKTA Start protein purification system to separate the cleaved and uncleaved proteins. The His-tag-cleaved GAF-A was recovered during the sample application step while the His-tagged TEV protease and residual uncleaved GAF-A were eluted with IMAC buffer B. The His-tag removed fractions were pooled, concentrated using Amicon Ultra-15 centrifugal unit (10 kDa MWCO), and buffer-exchanged into storage buffer. The purity of the proteins was confirmed by SDS-PAGE analysis before being aliquoted, flash-frozen, and stored in -80°C until usage.

**Expression and purification of DosS variants:**

The WT DosS and its E87N variant were expressed as per published protocols.[1] NWL DosS was expressed with the following modifications: the cells were grown to $OD_{600}$ of 0.6 before finally induced with 0.25 mM IPTG at 22°C for 24 hours.

DosS variants were purified following the GAF-A purification protocol with additional chromatographic methods. Specifically, the IMAC fractions containing full-length DosS were pooled, buffer-exchanged with ~90 mL DEAE buffer A (50 mM Tris-HCl, 1 mM EDTA, 5% glycerol, pH = 8.00) using an Amicon Ultra-15 centrifugal unit (50 kDa MWCO), before being subsequently applied onto a 5-mL HiTrap DEAE FF on AKTA Start system. After sample application, the column was washed with 7 CV of DEAE buffer A and the full-length DosS was eluted using a linear-gradient method: 0-100% of DEAE buffer B (50 mM Tris-HCl, 500 mM NaCl, 1 mM EDTA, 5% glycerol, pH = 8.00) for 12 CV. Fractions that corresponded to the peak of UV chromatogram were collected and the purity of the fractionated proteins was determined by SDS-PAGE. When further purification was necessary, the DEAE fractions were pooled, buffer-exchanged into storage buffer and loaded onto HiPrep 16/60 Sephacryl S-200 HR column. The protein was eluted at 0.15 mL/min using storage buffer. Subsequently, the fractions containing the DosS proteins were pooled, concentrated, flash-frozen, and stored in -80°C until usage.

**PCR and mutagenesis:**

Site-directed mutagenesis was performed in an Axygen Maxygene II Thermal Cycler (THERM-1000/THERM-1001) in Mode 1 (temperature control) using 0.2 ml flat cap PCR tubes (GeneMate) using Phusion Site-Directed Mutagenesis Kit (Thermo Fisher Scientific, Catalog no. F541) to obtain mutants. The primers used for E87N mutation were 5'-GGCGCCATGAATGTACATGAT-3' (forward) and 5'-ATCATGTACATTCAT GGCGCC-3' (reverse); for F98W mutation were 5'-GGTGTTACATTGGGTGTATGAGG-3' (forward) and 5'-CCTCATACACCCAATGTAACACC-3' (reverse) and for I121L mutation were 5'-GGGACTTGGGGTGCTCGGGTTGTTAATTG-3' (forward) and 5'-CAATTAACAACCCGAGCACCCCAAGTCCC-3' (reverse). The annealing temperatures which led to successful mutagenesis for primers E87N, F98W and I121L were 61 °C, 62 °C and 66.5 °C respectively. The PCR products were mixed with 6X TriTrackDNA loading dye (Thermo scientific, Ref no. R1161) and then loaded to a 0.8% agarose gel pre-stained with GelGreen nucleic acid stain (Biotium, Catalog no. 41104) along with GeneRuler 1 kb plus DNA ladder (Ref no. SM1331) to screen them. DNA gel electrophoresis was run on a Thermo scientific Owl Easycast B1A unit at 120 V for ~ 1.5 h to assess if the PCR was successful. Finally, NEB 5-alpha competent *E. coli* (High efficiency) cells were used for transforming the plasmids. After DNA purification through a GeneJET Plasmid Miniprep Kit (Thermo Fisher Scientific, Catalog no. K0502), the plasmids were sequenced with universal T7 promoter/terminator. DNA sequence was analyzed in BioEdit sequence alignment editor v. 7.2.1 and SnapGene viewer v. 5.0.7. DNA was quantified in a Nanodrop spectrophotometer using programmed methods.

**Characterization of proteins by gel electrophoresis:**

Purity of proteins was assessed by sodium dodecyl sulfate−polyacrylamide gel electrophoresis (SDS−PAGE) by running the protein samples through precast 4-12% Bis-Tris Plus Gels (Invitrogen Bolt) under denaturing conditions at room temperature in a Mini Gel Tank (Invitrogen, Catalog no. A25977. Running conditions were 200 V for 20-30 min. SeeBlue Plus2 Pre-stained protein standard (3-198 kDa, Invitrogen) or PageRuler Prestained protein ladder (10-170 kDa) was used as a marker. The images were scanned using a Typhoon FLA 9500 scanner (GE Healthcare) and analyzed through an ImageJ software.

**Hemochromogen assay:**

To determine the extinction coefficients of heme proteins in this work, a pyridine hemochromogen assay was performed according to the published protocol.[2] Briefly, 100 µL of ferric protein was mixed with 80 µL 5M NaOH and 800 µL pyridine. The fully oxidized spectrum was recorded before adding 10 uL of 0.5 M sodium dithionite solution to obtain the reduced mixture. The absorbance value at 557 nm was recorded and the molar absorptivity of pyridine-heme *b* hemochromogen (34.7$mM^{-1}cm^{-1}$) was used to finally determine the extinction coefficients of proteins.

**Crystallization of WT and mutant DosS GAF-A and structure determination:**

WT and mutant GAF-A DosS crystals were obtained with minor modifications to previously published methods.[3,4] Briefly, TEV-cleaved proteins were buffer exchanged into 20 mM Tris pH 7.5 using a PD-10 desalting column and concentrated to 7.7 mg/ml. Diffractable crystals were obtained utilizing hanging drop method and using seed stocks grown in ~200 mM calcium acetate,

100 mM Tris pH 7, 18% PEG 4000. The protein crystals were cryo-protected by soaking in the reservoir solution supplemented with ethylene glycol, by gradually increasing ethylene glycol concentration to 20%, and subsequently flash frozen in liquid nitrogen. X-ray diffraction data were collected at the NE-CAT beamline 24-ID-C of the Advanced Photon Source (Lemont, IL) and processed using XDS.[5] The structures were determined by molecular replacement with PHASER[6] using DosS GAF-A (PDB ID: 2W3D) as the search model. Iterative model building and refinement were conducted using COOT[7] and PHENIX[8]. A summary of crystallographic data statistics is shown in Table S2. Figures were generated using ChimeraX. The WT, F98W, and E87N GAF-A DosS structures have been deposited in the Protein Data Bank[9] with accession numbers of 9OTD, 9OTE, and 9OTF, respectively.

**<u>Measurement of $O_2$ affinity values for DosT and DosS proteins:</u>**

The $O_2$ affinity measurement and data analysis for DosT and DosS proteins were performed as described step-by-step in a methods paper published by our lab previously.[10]

**<u>MD studies with DosS GAF-A variants:</u>**

The crystal structure of GAF-A domains of DosS were used as the starting structure for MD simulations. The energy minimization, equilibration, and MD were performed on $O_2$-bound DosS GAF-A structures using methods similar to the ones described previously.[11]

**<u>NO binding studies with DosS variants:</u>**

Ferrous DosS variant was generated in the glovebag via reduction with 10x dithionite and removal of excess reductant with a PD-10 column and centricon. The reduced protein was diluted to ~ 2 μM in 50 mM Tris.HCl, 5% glycerol buffer and subjected to NO binding via addition of 5 equivalents of Proli NONOate. The mixture was incubate for ~ 10 minutes before taking the NO bound spectrum using CARY 60.

**<u>DFT studies with DosS GAF-A variants:</u>**

All calculations were performed using Gaussian 16 program. Initial wild type (WT) and the NWL active site models for DosS were based on DosS' X-ray structure of PDB file 2W3H and molecular dynamics simulation result, respectively. These models were subject to partial geometry optimization with protein residues truncated at Cα positions and the terminal $C_\alpha H_3$ groups fixed to mimic the protein environment on the basis of the previous studies of heme and non-heme iron proteins.[12–14] In the WT active site models, nearby residues F98, Y171, E87, I121, and P115 that could influence $O_2$ binding were included together with the proximal H149 and P146 residues. In contrast, the NWL active site models have the three corresponding F98W, E87N, and I121L mutations. In both WT and NWL models, the full heme cofactor is included, except that the propionic groups are substituted for propyl groups to avoid artificial H-bonding interactions as reported.[15] Geometry optimization and subsequent frequency calculations were performed using the hybrid HF-DFT method mPW1PW91 with the LanL2DZ basis for Fe, 6-311++G(2d,2p) basis for $O_2$, and five Fe-coordinated nitrogen atoms, and a 6-31G(d) basis for the rest atoms. This approach was selected on the basis of a methodological study[16] of several kinds of DFT methods including the more recently developed M06 and dispersion-corrected ωB97XD, on HNO and NO reactions via metal-containing systems.[17] Optimization calculations were conducted with the bulk protein environment simulated by the self-consistent reaction field method using the PCM approach[18] with a dielectric constant of 4.0 as previously reported. The subsequent frequency analysis was used to verify the nature of the stationary points on respective potential energy surfaces and to provide zero-point energy corrected electronic energies (EZPE's), enthalpies (H's), and Gibbs free energies (G's) at room temperature and 1 atm. Natural Population Analysis (NPA) charges and Mulliken spin densities were also calculated as implemented in Gaussian 16.

For WT and NWL $Fe^{II}$ systems, the singlet ground states for $O_2$ bound models were used. For $O_2$ bound protein models, both the open-shell singlet (OSS) ferric super oxy ($Fe^{3+}$-$O_2^{-}$) and closed-shell singlet (CSS) ferrous oxy ($Fe^{2+}$-$O_2$) structures were investigated with the lower energy OSS results used in the main text analysis. In addition, different conformations were studied to choose the most favorable one for discussion in the main text.

### Supplementary Tables and Figures:

**Table S1:** $O_2$ affinity constants ($K_d$) of heme-based oxygen sensors that feature tyrosine as the secondary coordination sphere (SCS) H-bonding residue

| Sensor | $O_2$ stabilizing SCS residue | Bacterial species | $K_d$ (μM) | Protein fold |
|---|---|---|---|---|
| H-NOX | Tyr140 | *C. subterraneous* | 0.048[19]<br>0.089[20]<br>0.023[10] | H-NOX |
| *Af*GcHK | Tyr45 | *Anaeromyxobacter sp. Fw109-5* | 0.077, 0.670[21] | GCS |
| YddV | Tyr43 | *E. coli* | 0.095, 14[22] | GCS |
| *Av*GReg | Tyr44 | *Azotobacter vinelandii* | 0.120[23] | GCS |
| DosS | Tyr171 | *M. tuberculosis* | 3[24]<br>0.58[25]<br>0.46[1] | GAF |
| *Bpe*GReg | Tyr43 | *Bordetella pertussis* | 0.64[26] | GCS |
| HemAT | Tyr70, Thr95 | *Bacillus subtilis* | 4.5,100[27] | GCS |
| DosT | Tyr169 | *M. tuberculosis* | 26[24]<br>3.3[1] | GAF |
| H-NOX | Tyr139 | *C. botulinum* | 53[28] | H-NOX |

**Table S2: Crystallization data collection and refinement statistics**

| | DosS-WT | F98W | E87N |
|---|---|---|---|
| **Data collection** | | | |
| Space group | $P2_12_12_1$ | $P2_12_12_1$ | $P2_12_12_1$ |
| Unit cell dimensions | | | |
| *a, b, c (Å)* | 33.14, 77.40, 110.10 | 36.48, 75.85, 112.74 | 33.140, 75.69, 110.65 |
| α, β, γ (°) | | | |
| Resolution (Å) | 33.16 - 2.06 (2.22 - 2.06) | 56.37 - 1.90 (1.97 - 1.90) | 44.66 - 2.29 (2.52 - 2.29) |
| *Rsym or Rmerge* | 0.129 (1.341) | 0.039 (1.574) | 0.068 (1.317) |
| *I / σI* | 7.9 (1.3) | 14.9 (1.0) | 12.1 (1.4) |
| Completeness (%) | 91.5 (52.0)* | 99.2 (97.7) | 90.9 (46.0)* |
| Redundancy | 7.2 (6.7) | 5.0 (5.0) | 7.3 (7.8) |
| CC1/2 | 0.995 (9.531) | 1 (0.999) | 0.999 (0.623) |
| **Refinement** | | | |
| Resolution (Å) | 33.16 - 2.06 (2.22 - 2.06) | 56.37 - 1.90 (1.97 - 1.90) | 44.66 - 2.29 (2.52 - 2.29) |
| No. Reflections | 14129 (732) | 25157 (1281) | 11879 (611) |
| *Rwork / Rfree* | 0.218 / 0.264 | 0.207 / 0.243 | 0.229 / 0.269 |
| No. atoms | 2292 | 2385 | 2282 |
| Protein | 2191 | 2251 | 2185 |
| Ligand/ion | 90 | 105 | 88 |
| Water | 11 | 29 | 9 |
| *B-factor* | 62.12 | 70.58 | 89.57 |
| Protein | 62.92 | 71.22 | 90.91 |
| Ligand/ion | 44.53 | 58.34 | 59.23 |
| Water | 47.44 | 65.02 | 62.34 |
| Ramachandran plot | | | |
| Favored (%) | 95.71 | 96.48 | 92.81 |
| Allowed (%) | 4.29 | 3.52 | 6.47 |
| Outliers (%) | 0 | 0 | 0.72 |
| R.m.s. deviations | | | |
| Bond lengths (Å) | 0.003 | 0.007 | 0.003 |
| Bond angles (°) | 0.68 | 0.96 | 0.69 |

Statistics for the highest-resolution shell are shown in parentheses.

*Ellipsoidal

| | | | |
|---|---|---|---|
| a*, b*, c* (Å) | 2.452, 2.082, 2.036 | | 2.301, 2.235, 2.296 |

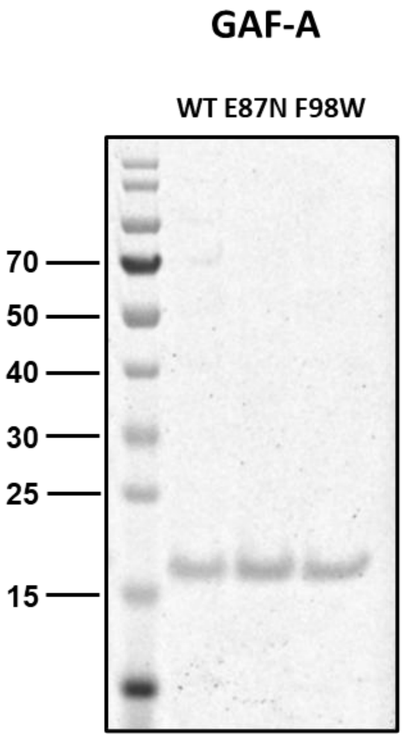


**Fig. S1:** SDS PAGE gel image demonstrating WT, E87N and F98W GAF-A DosS. Expected MW: WT DosS GAF-A = 16,319 Da; E87N DosS GAF-A = 16,305 Da; F98W DosS GAF-A = 16,359 Da.

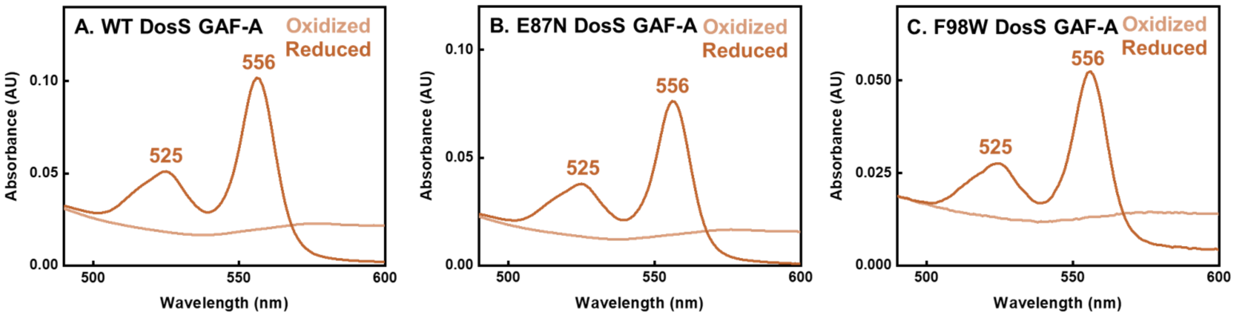


**Fig. S2**: Pyridine hemochromogen assays were performed on (a) WT DosS, (b) E87N, and (c) F98W GAF-A DosS to ascertain the presence of heme *b* and measure the extinction coefficients of proteins based on the spectral features. Spectra of reduced and oxidized pyridine hemochromogen with notable peaks shown. The extinction coefficients were calculated as 160 ± 6 $mM^{-1}$ $cm^{-1}$ (n = 3), 157 ± 8 $mM^{-1}$ $cm^{-1}$ (n = 3), and 110 ±1 $mM^{-1}$ $cm^{-1}$ (n = 3) for WT, E87N, and F98W GAF-A DosS respectively.

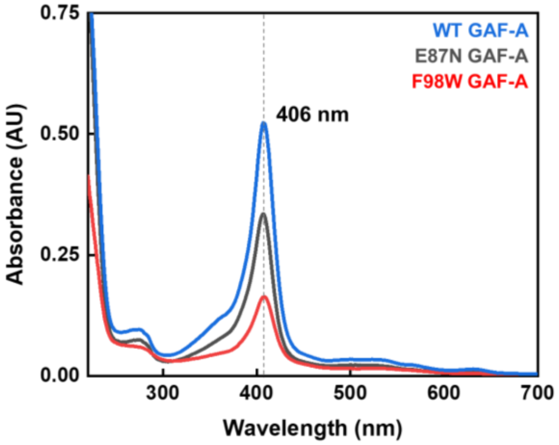


**Fig. S3**: UV-Vis spectra of ferric forms of WT, E87N and F98W GAF-A DosS

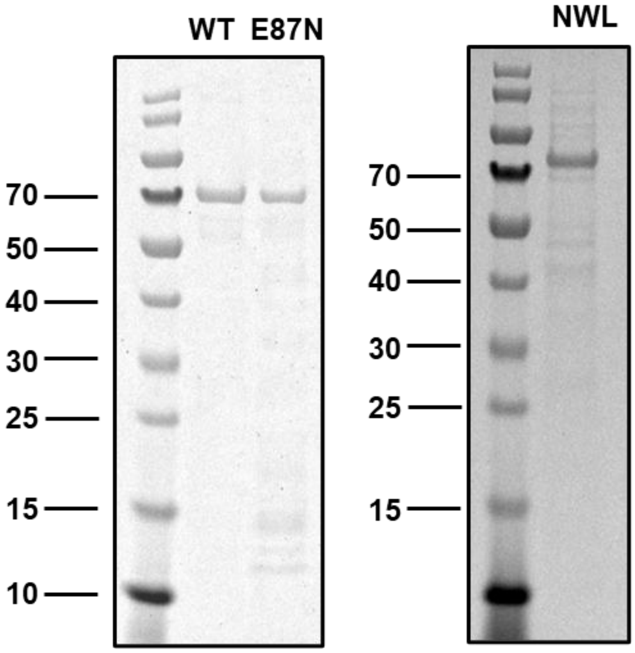


**Fig. S4:** SDS gel demonstrating WT, E87N and NWL DosS. Expected MW: WT DosS = 64,047 Da; E87N DosS = 64,031 Da; NWL DosS = 64,070 Da.

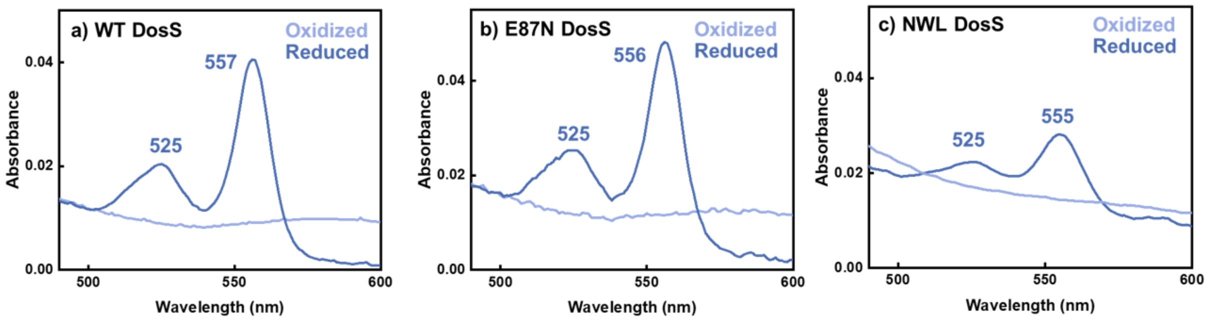


**Fig. S5**: Pyridine hemochromogen assays were performed on (A) WT DosS, (B) E87N DosS, and (C) NWL DosS to ascertain the presence of heme *b* and find the extinction coefficients of the proteins based on the spectral features. Spectra of reduced (dark blue) and oxidized (purple) pyridine hemochromogen with notable peaks are shown. The extinction coefficients were calculated as 145 ± 2 $mM^{-1}$ $cm^{-1}$ (n = 3), 164 ±6 $mM^{-1}$ $cm^{-1}$ (n = 3), and 82 ± 7 $mM^{-1}$ $cm^{-1}$ (n = 2) for WT, E87N, and NWL DosS (oxy-form), respectively. Note that NWL DosS spectra have undergone adjacent averaging smoothing signaling process in Origin software to reduce background noises.

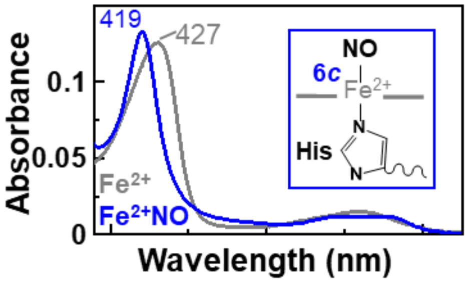


**Fig. S6**: UV-Vis spectra showing ferrous E87N DosS (grey) binding NO to form 6c ferrous-NO complex (blue).

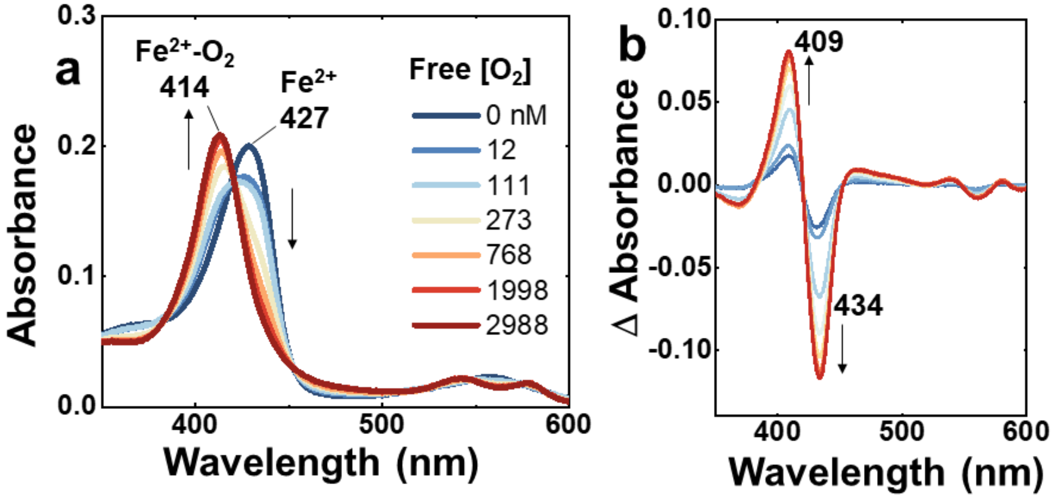


**Fig. S7:** UV–Vis spectral changes in E87N DosS upon binding $O_2$ at various free $O_2$ concentrations measured using the optode. c) Difference spectra showing spectral changes when E87N DosS binds $O_2$.

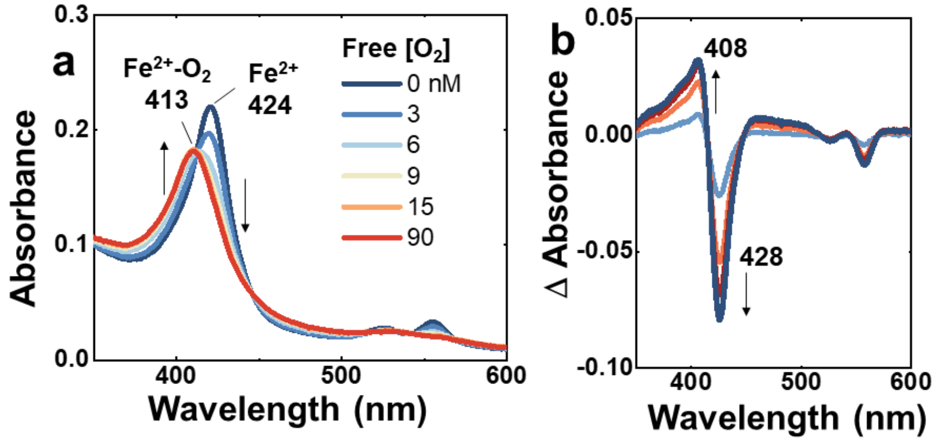


**Fig. S8:** UV–Vis spectral changes in NWL DosS upon binding $O_2$ at various free $O_2$ concentrations measured using the optode. c) Difference spectra showing spectral changes when DosT binds $O_2$.
